# Large-scale meta-analysis suggests low regional modularity in lateral frontal cortex

**DOI:** 10.1101/083352

**Authors:** Alejandro de la Vega, Tal Yarkoni, Tor D. Wager, Marie T. Banich

## Abstract

Extensive fMRI study of human lateral frontal cortex (LFC) has yet to yield a consensus mapping between discrete anatomy and psychological states, partly due to the difficulty of inferring mental states in individual studies. Here, we used a data-driven approach to generate a comprehensive functional-anatomical mapping of LFC from 11,406 neuroimaging studies. We identified putatively separable LFC regions on the basis of whole-brain co-activation, revealing 14 clusters organized into three whole-brain networks. Next, we used multivariate classification to identify the psychological states that best predicted activity in each sub-region, resulting in preferential psychological profiles. We observed large functional differences between networks, suggesting brain networks support distinct modes of processing. Within each network, however, we observed low functional specificity, suggesting discrete psychological states are not modularly organized. Our results are consistent with the view that individual LFC regions work as part of highly parallel, distributed networks to give rise to flexible, adaptive behavior.

---

Decades of research have suggested lateral frontal cortex (LFC) plays a critical role in the execution of flexible, goal-directed behavior^1^. Such flexible behavior enables the navigation of complex, rapidly changing environments, the pursuit of distant goals in the face of various obstacles, planning for hypothetical future events, and the communication of complex ideas with others using language. Although extensive work has identified putatively separable psychological processes critical for flexible behavior^2^– such as ‘working memory’, ‘inhibition’ and ‘conflict’– the precise organization of these processes within discrete lateral frontal anatomy remains actively debated.

Much progress has been made in understanding the LFC’s functional organization by identifying putatively separable LFC subregions on the basis of properties that constrain information processing. For instance, discrete regions have been proposed based on differences in anatomical microstructural properties (e.g. cytoarchitecture^3^), and anatomical^4–6^ and resting-state functional connectivity^7,8^. Although these studies have helped carefully characterize important functional properties of LFC, it is unclear to what extent the boundaries derived from such methods correspond to the organization of brain activity observed during distinct psychological states^9^.

One approach used to map the functional correlates of distinct behavioral phenotypes is the quantitative meta-analysis of functional MRI (fMRI) studies. Such meta-analyses help overcome the low power observed in individual fMRI studies and produce more precise spatial maps of psychological states that activate LFC, such as working-memory^10,11^, inhibition^12^, switching^13,14^, language^15^, mentalizing^16^ and self-referential processing^17^. However, due to the effort required to compile meta-analyses, and because most researchers are interested in a particular psychological domain, most meta-analyses are typically focused on a particular subregion of LFC or a subset of domain-specific set of psychological processes.

The narrow scope of most existing meta-analyses necessarily limits the extent of their impact for two reasons. First, complex behavior likely results from the coordinated activity of individual regions participating across whole-brain networks^18^; thus, it is critical to interpret the function of each region in a broader context in order to understand their role within large-scale networks and to better identify subtle differences between similar regions in the same network. Second, it is notoriously difficult to infer mental function from observed brain activity (the so-called problem of “reverse inference”^19^), as determining the relative specificity with which a particular task or process activates a given region requires the ability to quantify the likelihood of activation in that region across a wide range of potential tasks. This problem is particularly acute in brain regions that appear to activate frequently across a broad range of tasks. Hence, the fact that LFC appears to be involved in a broad range of tasks—putatively due to its critical role in guiding flexible behavior^1,20,21^—implies that subregions of this area may be particularly difficult to associate with specific mental operations^22^.

Here we address these issues by creating a comprehensive mapping between data-derived semantic topics representing psychological states and LFC using Neurosynth^23^, a framework for large-scale fMRI meta-analysis composed of nearly 11,500 studies. First, we used a data-driven method that exploits the observation that functionally related regions co-activate across studies^24–28^ to cluster individual voxels into putatively separable subregions (Figure 1a). We applied clustering at two spatial scales, identifying three distinct whole brain networks in LFC composed of several smaller subregions with dissociable co-activation patterns (Figure 1b). We then characterized the functional profile of each resulting region using multivariate classification, contrasting studies that activated each region with those that did not, revealing dissociable psychological profiles for each LFC subregion (Figure 1c). Collectively, we provide a comprehensive and relatively unbiased meta-analytic functional-anatomical mapping of LFC.

**Figure 1.**
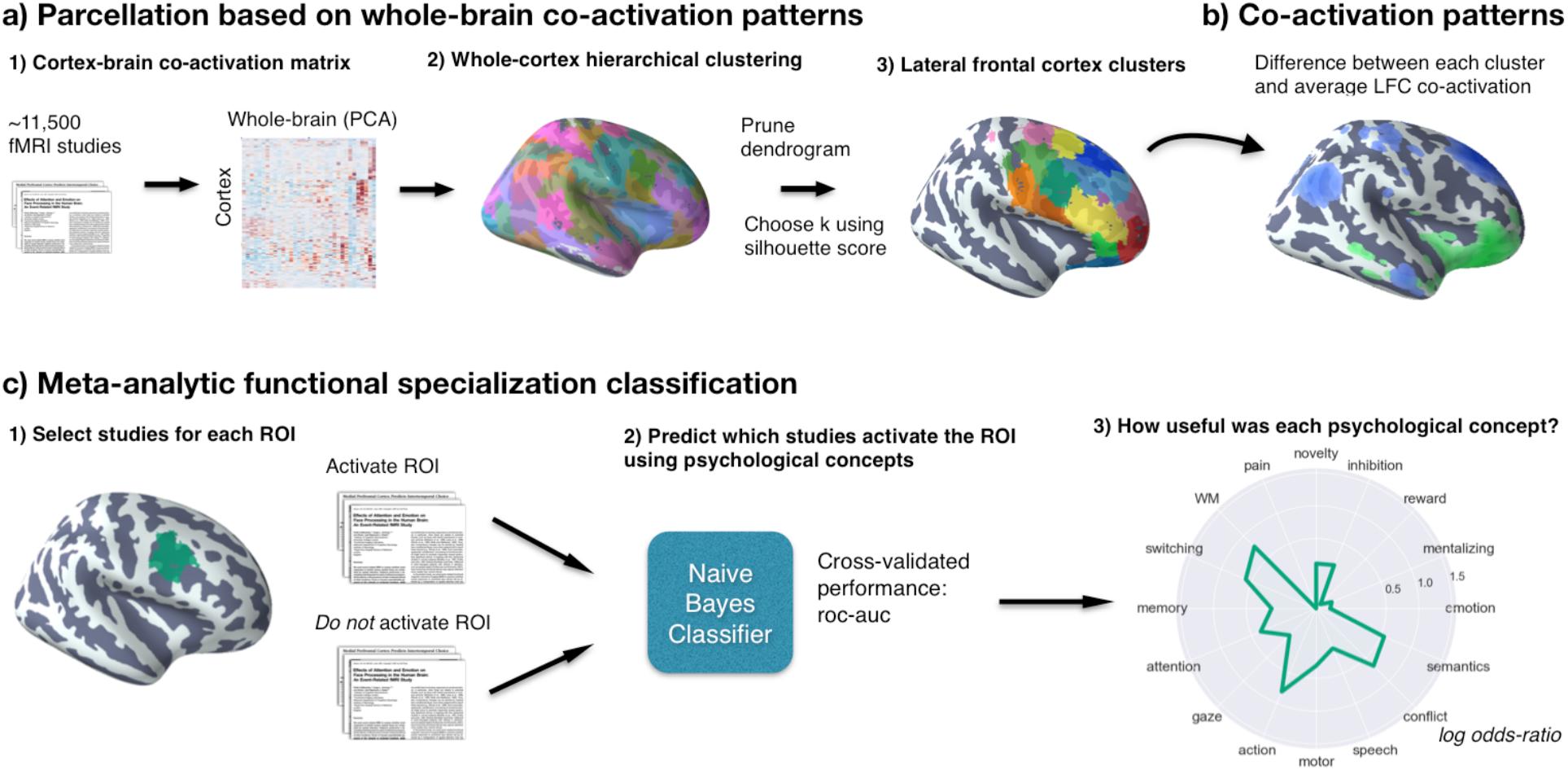
Methods overview. a) We calculated co-activation across studies between every cortical voxel and the rest of the brain, including subcortex. We then applied Ward hierarchical clustering to obtain whole-brain clustering results. We chose two spatial scales to focus on using the silhouette method^27,29^ and selected clusters in LFC from the whole-brain clustering solutions. b) We contrasted the whole-brain co-activation of each cluster with LFC at large, identifying voxels across the brain that showed differential co-activation. c) We generated functional preference profiles for each cluster by determining which latent psychological topics^30^ best predicted the cluster’s activation across studies in the database.

## Results

### Hierarchical clustering of lateral frontal cortex

We identified spatially dissociable regions on the basis of shared co-activation profiles with the rest of the brain^24–28^, an approach that exploits the likelihood of a voxel co-activating with other voxels across studies in the meta-analytic database. To identify whole-brain networks spanning beyond LFC, we applied hierarchical clustering to the whole cortex and selected clusters within LFC mask for further analysis (Figure 2b). In order to map structure to function across various spatial scales, we extracted 4– to 100– whole-brain clusters and evaluated their quality using the silhouette score, a measure of intra-cluster cohesion^27,29^ (Figure 2a). Given the intractable nature of choosing the ‘correct’ number of clusters^31^ and the lack of a single dominant solution in our data, we focused on two well spaced granularities, 5 and 70 whole-brain clusters, avoiding low quality solutions (i.e. 7-38 clusters). Importantly, we do not argue that the solutions we selected are in any way privileged, nor did we aim to match the scale of previous parcellations; rather, we simply chose two spatial scales for subsequent analysis with distinct vantage points into the hierarchical organization of LFC.

**Figure 2.**
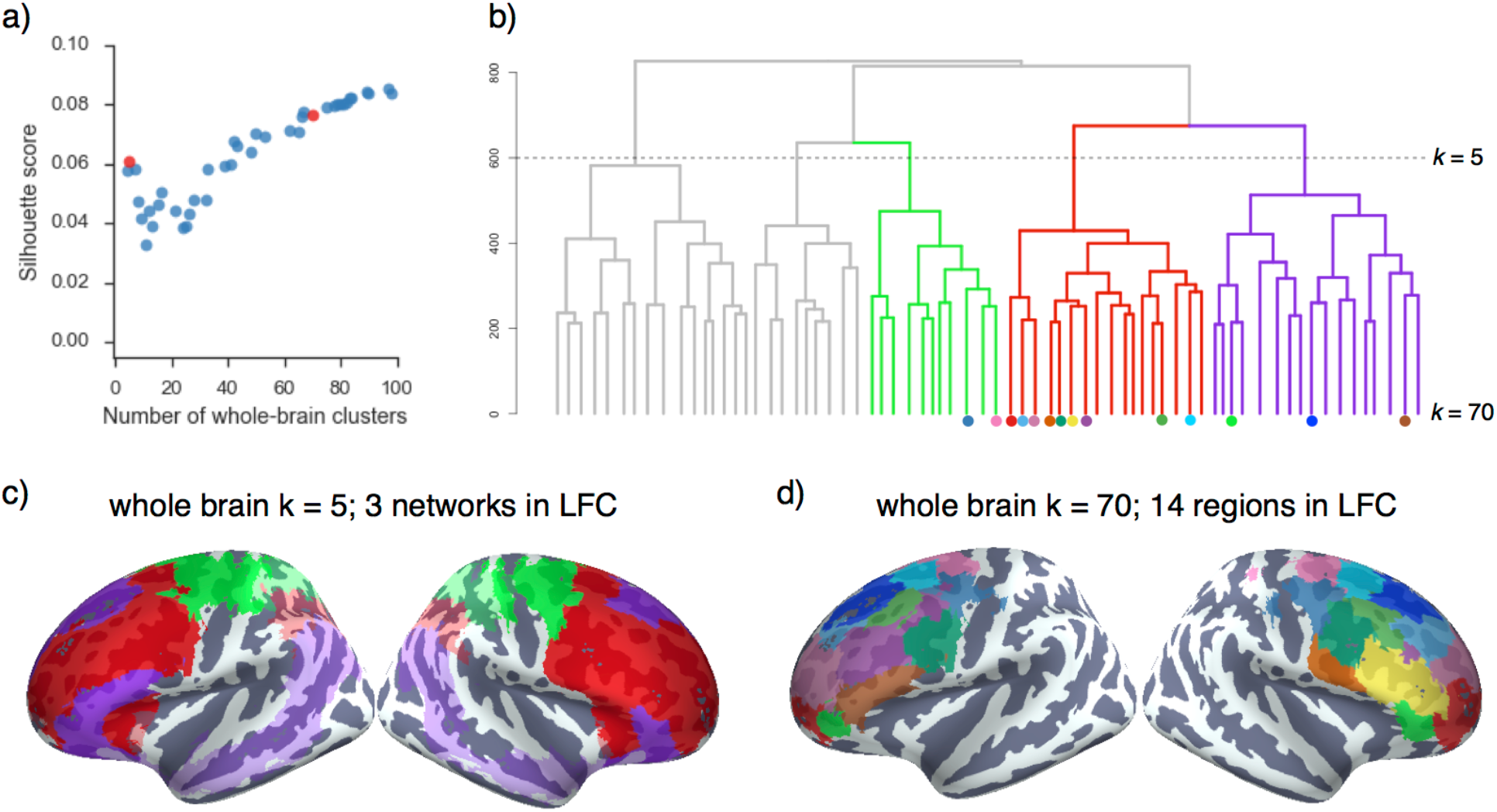
Whole-cortex co-activation based hierarchical clustering reveals 3 networks in lateral cluster that fractionate into constituent subregions. a) The silhouette score, a measure of intra-cluster cohesion, was used to select two spatial scales: 5 and 70 whole-brain clusters. b) Whole brain hierarchical clustering dendrogram. Color-coded branches correspond to three of five whole-brain networks in LFC and color-coded nodes correspond to 14 LFC regions from 70 whole-brain clusters. c) Clusters at *k =* 5 revealed three clusters in LFC resembling large-scale brain networks: “fronto-parietal” (red), “default” (purple) and “somatosensory-motor” (green) d) Clusters at *k =* 70 revealed 14 clusters with a 75% of their voxels in LFC.

To understand the large-scale network organization of LFC, we focus on the five-cluster solution as this scale exhibited the greatest silhouette score of coarse network-level solutions (see SI Figure 1 for whole-brain cluster results). Three of these whole-brain network clusters were present in LFC (Figure 2c) and showed moderate correspondence to previously described large-scale networks^32^. Although these clusters were not isomorphic with resting-state networks^32,33^, these results are consistent with the view that large-scale brain networks supersede gross anatomical boundaries, such as LFC, as functional-organizational units.

The largest of the three clusters, which we refer to as the “fronto-parietal” network, spanned half of LFC, primarily in prefrontal cortex, and resembled Yeo et al., 2011’s description of the “fronto-parietal” network^32^ (dice coefficient (d) = 0.56). Additionally, this cluster spanned medial-frontal and anterior insular aspects of the “ventral attention” network (d = 0.21) (See SI Figure 2 for a cross reference between our networks and Yeo et al., 2011). A second cluster, which we refer to as the “default” network, closely matched extensive descriptions of the “default” or “task negative” network (d = 0.62)^34^. The final cluster, which we refer to as the “sensorimotor” network, was located in posterior LFC and showed moderate overlap with Yeo’s “somatosensory-motor” network (d = 0.36) and, to a lesser extent, the “dorsal attention network” (d=0.31).

Having identified large-scale networks in LFC, we sough to identify more functionally specific subregions within each network with potentially dissociable psychological profiles. Although the silhouette values indicated that inter-cluster cohesion continuously increases with number of clusters, we chose to focus on a spatial scale that balanced clustering quality with psychological interpretability. Thus, we chose to focus on the 70- cluster solution, as this was the coarsest scale to result in a set of largely spatially contiguous LFC clusters. From these 70 whole brain clusters, we identified 14 clusters within our LFC mask (Figure 2d), hierarchically organized into the coarser large-scale networks.

To provide direct insight into the functions of the 14 LFC fine-grained clusters we identified, we applied two approaches. First, we determined which voxels across the brain differentially co-activated with each cluster, revealing distinct patterns of whole brain co-activation. Second, we used semantic data from Neurosynth to determine which latent psychological topics predict the activation of each cluster, resulting in a meta-analytic psychological preference profile for each subregion. Next, we step through these results separately for each network.

### Fronto-parietal network

The majority of lateral frontal cortex belonged to the frontal extent of the “fronto-parietal” network, which further spanned portions of lateral parietal cortex, anterior insula, pre-SMA, mid-cingulate cortex (MCC), and the precuneus. Within LFC, we identified 10 finer-grained subregions within the “fronto-parietal” network. For purely illustrative purposes, we used the hierarchical clustering dendrogram (Figure 2b) to identify a granularity in which these clusters formed three sets; at *k =* 24 whole brain clusters, these 10 LFC clusters organized into three groups: caudal, mid and rostral regions. Across these three groupings, all clusters showed robust associations with executive functions, although we observed subtle variations in psychological preferences.

In caudal LPFC, we identified two adjacent bilateral clusters (Figure 3a). The most posterior of the two (‘6/8’) was located anterior to the premotor cortex and extended from lateral superior frontal gyrus to the intermediate frontal sulcus of middle frontal gyrus. This cluster overlapped with functional descriptions of the frontal eye fields (FEF)– a region important for volitional eye saccades^35^.

Immediately anterior, we identified a cluster (‘9/46c’) spanning caudal area 9/46 from the intermediate frontal sulcus into caudal portions of 9/46v. Notably, although cluster ‘9/46c’ arguably extends well into “mid” LPFC, this cluster did not group with other mid-LPFC clusters until much coarser granularities, suggesting these clusters may exhibit distinct functional signatures despite their spatial proximity.

Anterior and ventral to caudal LPFC, we identified four clusters spanning common definitions of ‘mid’ lateral prefrontal cortex (Figure 3b). The organization of clusters in this region, however, varied by hemisphere. Most dorsally, we identified a mostly left-lateralized cluster (‘9/46v’), extending from the intermediate frontal sulcus into the fundus of the inferior frontal sulcus. Next, we identified a cluster, which we refer to as right IFG (‘IFG [R]’), spanning the majority of area BA45 in the right hemisphere. Notably, only right IFG was part of the fronto-parietal network, consistent with the observation that this region is consistently observed during goal-directed cognition. Posterior to these two clusters, we identified a bilateral cluster consistent with the inferior frontal junction (‘IFJ’) (e.g. MNI coordinates: 48, 4, 33^36^), located in the fundus of caudal inferior frontal sulcus, extending into precentral, inferior frontal and middle frontal gyri. Finally, ventral to this cluster, but only in the right hemisphere, we identified a fourth cluster (‘44 [R]’) located in posterior IFG, spanning BA44 and abutting BA6.

**Figure 3.**
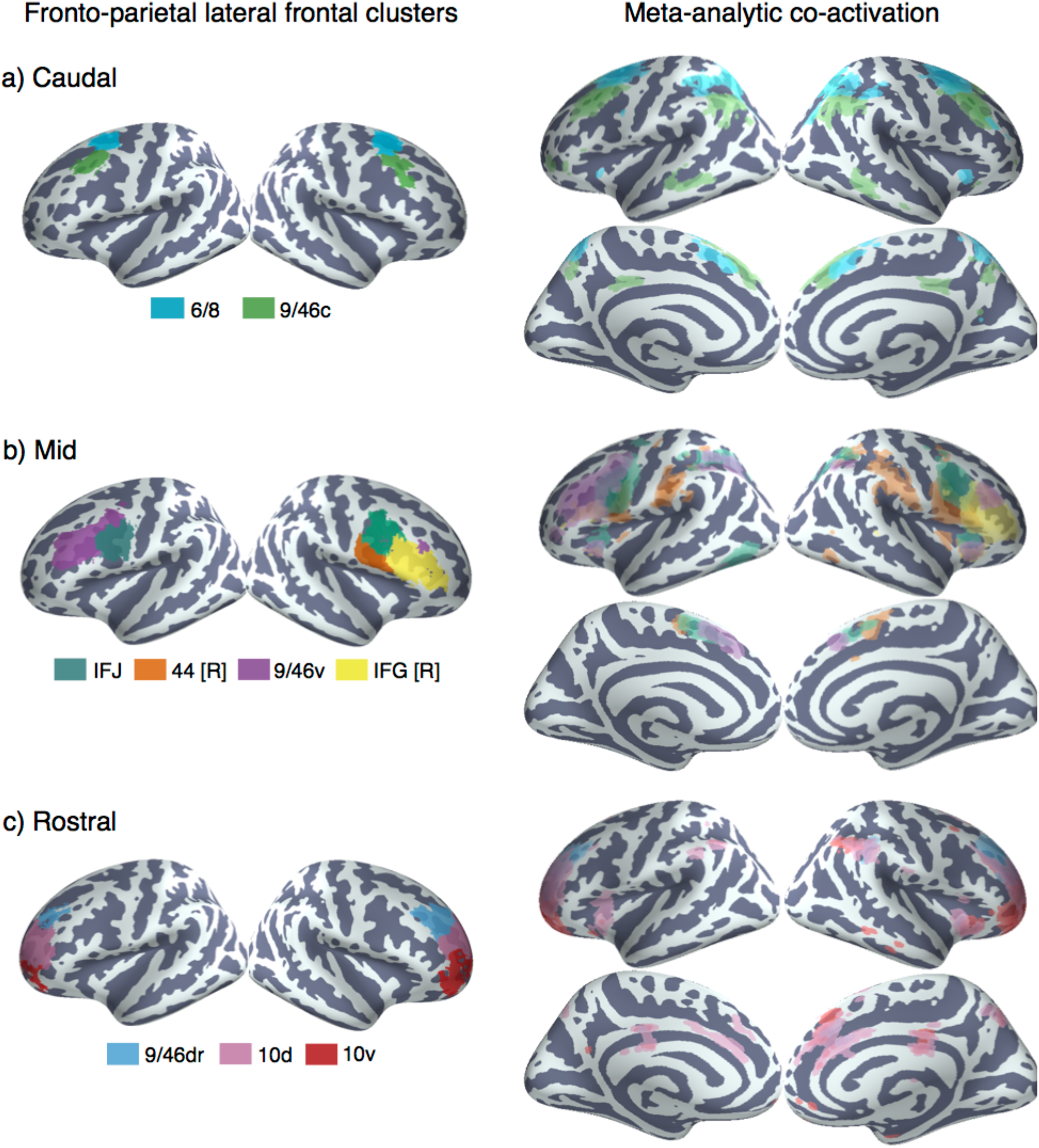
Anatomical location and meta-analytic contrast of lateral frontal clusters of the fronto-parietal network. Left: a) Two clusters located in caudal frontal cortex. b) Four clusters located in mid-lateral pre-frontal cortex. c) Three clusters located in rostral lateral pre-frontal cortex. Clusters were assigned labels corresponding to cytoarchitechtonic areas^3^ whenever possible. In cases where the region spanned multiple cytoarchitechtonic areas, broader anatomical (e.g. inferior frontal junction [IFJ]) labels were assigned. Right: Meta-analytic co-activation contrast of fronto-parietal LFC. Colored voxels indicate significantly greater co-activation with the seed region of the same color than other lateral frontal regions in the fronto-parietal network. Images are presented using neurological convention and are corrected using false discovery rate (FDR; q = 0.01).

In ‘rostral’ LPFC, we identified three bilateral clusters spanning BA10 (Figure 3c). These three clusters were organized along a ventral-dorsal axis, consistent with a prior DTI parcellation^4^, and were exclusively in lateral frontal cortex, consistent with cytoarchitechtonic evidence of a lateral-medial distinction of the frontal pole^37^. The most dorsal cluster (‘9/46dr’) extended into rostral portions of BA 9/46, while the next two clusters (‘10v’ and ‘10D’) were exclusively located in BA 10, separated along a dorsal/ventral axis

#### Meta-analytic co-activation

To better understand functional differences between these regions, we directly contrasted the co-activation of each cluster with that of LFC as whole in order to identify voxels across the brain that differentially co-activated with each cluster (Figure 3; right panel). Strikingly, we observed that most differential co-activation occurred within other cortical association cortex areas such as lateral parietal cortex (LPC), pre-SMA and MCC, and the insula. Across LPC, each LFC cluster co-activated most strongly with distinct areas across a gradient extending from tempo-parietal junction to the lateral parieto-occipital sulcus. For example, clusters ‘9/46c’ and all fronto-polar clusters showed greater co-activation with parietal cortex ventral to the intraparietal sulcus. In contrast, area ‘6/8’ and all four ‘mid’ LPFC clusters showed greater co-activation within and dorsal to the intraparietal sulcus.

Similarly in medial PFC, all clusters except right IFG and ‘9/46dr’ co-activated most strongly with slightly different portions of pre-SMA and MCC. Generally, more anterior clusters co-activated more strongly with more anterior portions of pre-SMA/MCC. For instance, ‘10d’ co-activated most strongly with a anterior mid-cingulate cortex while ‘44 [R]’ co-activated most strongly with the SMA. Finally, in the insula, several LFC subregions exhibited differential co-activation with distinct sub-divisions of the insula. For example, cluster ‘44 [R]’ co-activated most strongly with the posterior insula– an important region for pain and sensorimotor processing^38^– whereas IFJ co-activated most strongly with the dorsal anterior insula, a subregion implicated in goal-directed cognition. In contrast, area 10v showed greater co-activation with ventral anterior insula, an area implicated in affect^38^.

This observation that the bulk of co-activation differences between LFC subregions of the fronto-parietal network occurred within other cortical association areas is consistent with the hypothesis that association cortex is composed of parallel interdigitated networks^32^. That is, these findings suggest subregions of the FPN do not participate with categorically distinct sets of regions across the brain, and instead perform subtly different roles within a distributed network.

#### Meta-analytic functional preference

Next, we used a data-driven approach that surveyed a broad range of fMRI studies to quantify the degree to which distinct psychological states might be preferentially associated with different LFC clusters (Figure 1c). We trained naïve Bayes classifiers to predict the presence or absence of activation in each LFC cluster using a set of 60 psychological topics derived by applying a standard topic modeling approach to the abstracts of articles in the Neurosynth database^39^. We used the fitted model coefficients to quantify the strength of association between each psychological topic and the presence of activation in the corresponding LFC cluster (measured as the log odds-ratio [LOR] of the probability of each topic in studies that activated a given cluster relative to the probability of the same topic in studies that did *not* activate the cluster). Values greater than 0 indicate that the presence of that topic in a study positively predicts activity in a given region. We report the results of 16 psychological topics that loaded strongly onto LFC regions (Table 1) and restrict interpretation to significant associations using False Discovery Rate (FDR; q < 0.01). In addition, whenever we comparatively discuss sets of regions, we discussed differences if the 95% confidence interval (CI) of a given topic did not overlap between two regions (SI Figure 3). As the latter comparisons are post-hoc and exploratory, caution in interpretation is warranted.

Consistent with a distributed role for the fronto-parietal network in goal-directed cognition, all nine clusters were significantly associated with working-memory, all clusters except 10d and 10v were associated with conflict, and seven clusters were associated with switching (Figure 4). The present results are inconsistent with focal anatomical locations for high-level executive processes; instead, these results suggest that distributed activation across fronto-parietal network supports goal-directed cognition in the face of interference and conflict^40^. Despite the relatively low modularity we observed across the fronto-parietal network, multivariate associations between individual subregions and psychological states suggest preferential functional correlates can be identified for each cluster.

#### Caudal fronto-parietal LFC

Consistent with its co-location with the frontal eye fields, ‘6/8’ was the only cluster significantly associated with saccadic eye movements (i.e. ‘gaze’) in the fronto-parietal network, and was also associated with ‘attention’. This pattern suggests that area ‘6/8’ may be important for directing attention to relevant external stimuli to support downstream information processing. However, ‘6/8’ was also significantly associated with a ‘working-memory’ topic, consistent with a recent lesion study implicating the FEF in a causal role in working memory^41^. These results suggest area ‘6/8’ is not merely involved in low-level saccadic eye movements, but plays an important role in higher-level cognition.

In contrast, cluster ‘9/46c’ showed a much less distinctive functional signature, with relatively weak associations to other psychological processes outside of core EF processes and ‘memory’. This relatively diffuse pattern suggests area ‘9/46c’ may be involved in domain-general processes that span across distinct psychological states in our topic model.

**Figure 4.**
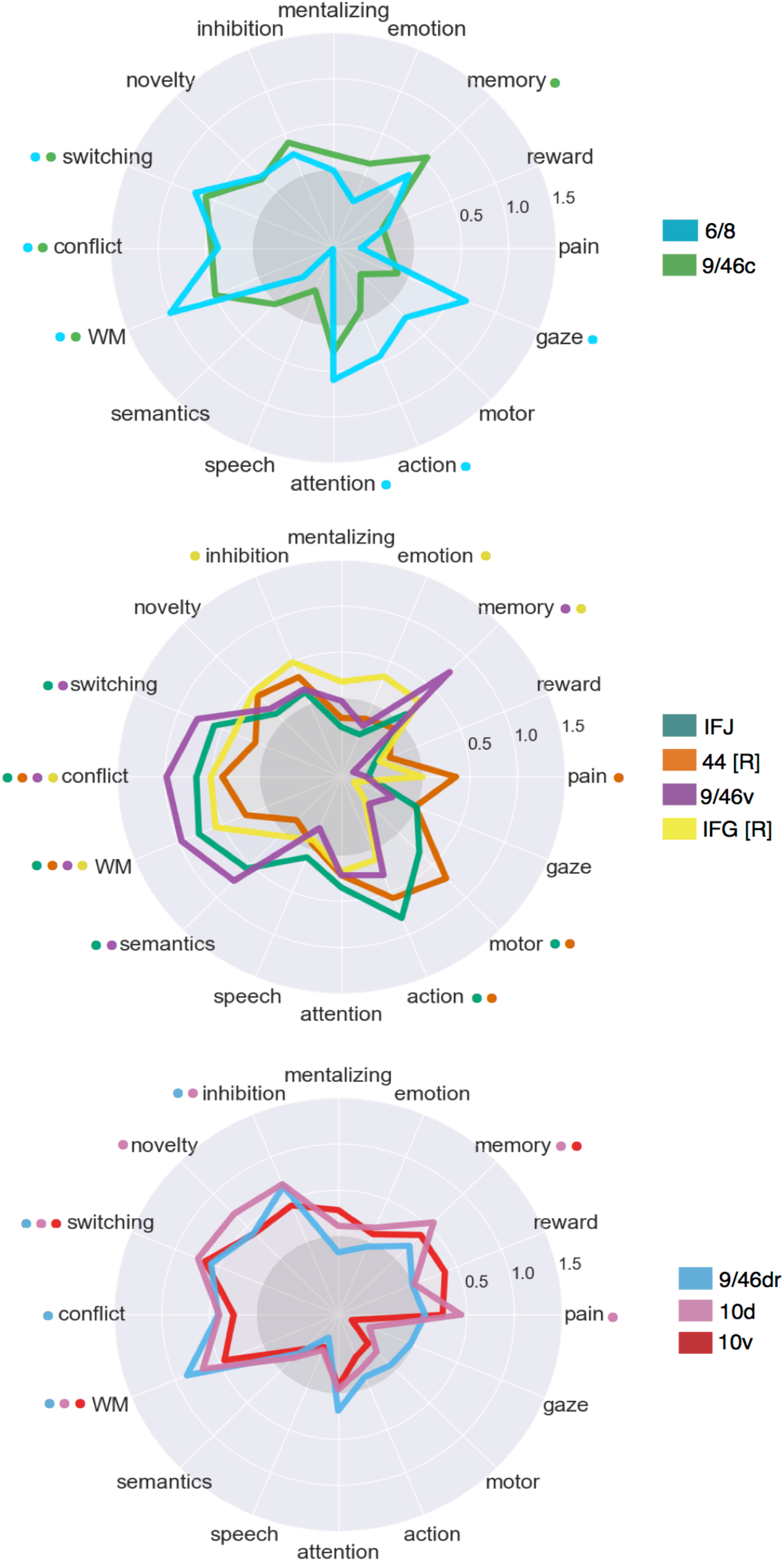
Meta-analytic functional preference profiles for lateral frontal regions in the fronto-parietal network. Each cluster was profiled to determine which psychological topics best predicted its activation. Each of the three functional groups we identified showed distinct functional profiles, although appreciable variation was observed for each individual cluster.

Strength of association is measured in log odds-ratio (LOR), and permutation-based significance corrected using false discovery rate (FDR) of q = 0.01 is indicated next to each psychological concept by color-coded dots corresponding to each region. Negative associations are indicated by the grey circle.

#### Mid fronto-parietal LFC

Clusters ‘9/46v’ and IFJ showed similar functional profiles, exhibiting robust associations with several executive functions (e.g. ‘working memory, ‘conflict’, ‘switching’) in addition to ‘semantics’. Cluster ‘9/46v’ showed a particularly strong association with executive functions– exhibiting the strongest relationship across LFC with ‘conflict’; these results are consistent with a hypothesized role for mid-DLPFC as a seat of high-level executive processes^3^.

These results are also consistent with the hypothesis that IFJ is involved in task-set switching; ^13^ however, given that several other LFC clusters were similarly associated with switching, it is unlikely IFJ is focally responsible for this phenomenon. Yet, IFJ showed a stronger association with low and high level motor function (i.e. ‘motor’, ‘action’) than other fronto-parietal LFC clusters, suggesting that IFJ is important for motoric aspects of cognitive control^42^. In contrast, cluster ‘44 [R]’ exhibited weaker associations with executive functions and robust associations with motor function and ‘pain’, suggesting area this area is more involved in sensori-motor processing than high-level cognitive control.

Finally, ‘IFG [R]’ showed a distinct functional signature to other mid LPFC clusters, with much weaker associations with conflict, working memory and switching; instead, ‘IFG [R]’ was associated with ‘inhibition’– consistent with extensive studies linking this region to inhibitory processes^10^. ‘IFG [R]’ was also associated with ‘emotion’, consistent with the hypothesis that this region is crucial for effective emotion regulation and reappraisal^43,44^. However, the relationship between ‘inhibition’ and ‘IFG [R]’ was not particularly strong or significantly greater than other fronto-parietal regions, suggesting ‘IFG [R]’ may play a more domain general role such as context monitoring^45^. Alternatively, local neuronal groups not detectable by fMRI may play more specific and distinct roles^46–48^.

#### Rostral fronto-parietal LFC

Although ‘rostral’ fronto-parietal clusters exhibited significant associations with various executive processes, these three clusters were characterized by weaker associations with motor function (‘action’), language and ‘conflict’. Rather, clusters ‘9/46dr’ and ‘10d’ were robustly associated with ‘inhibition’ and cluster ‘10d’ with ‘novelty’. This pattern of results suggests fronto-polar LFC regions may be important for high-level monitoring and guiding of cognitive control, removed from low-level motor implementations.

Finally, the most ventral fronto-polar region, cluster ‘10v’, showed a more distinct pattern, exhibiting weaker associations with all executive processes but a significant association with ‘reward’ (at a lower threshold, q<0.05). This pattern is consistent with its location near orbitofrontal cluster, and provides support for hypotheses that suggest that the ventral frontal pole may be important for representing the value of appetitive stimuli in order to effectively guide goal-directed behavior^4^.

### Default network

#### Anatomical correspondence

We identified three distinct default network clusters in LFC, consistent with previous descriptions of the default network and large-scale rs-fMRI parcellations (Figure 5a)^32^. The first two clusters were positioned adjacent to each other in ventrolateral prefrontal cortex. The more anterior of the two (‘47/12’) spanned lateral orbitofrontal cortex and IFG orbitalis bilaterally, while a more posterior and dorsal cluster spanned inferior frontal gyrus exclusively in the left hemisphere (‘IFG [L]’). Finally, we identified a third cluster in dorsal LPFC consistent with BA9^3^, extending from superior frontal gyrus to dorsal middle frontal gyrus across the superior frontal sulcus. This cluster has long been noted for its lack of anatomical input from lateral and medial parietal cortex^49,50^. Thus, despite these cluster’s close proximity to fronto-parietal clusters, we expected them to exhibit very distinct functional profiles.

#### Meta-analytic co-activation

Consistent with the grouping of these clusters with the default network, clusters ‘47/12’ and ‘9’ co-activated much more strongly than the rest of LFC with other default network regions, such as dorsal medial PFC (mPFC), middle temporal gyrus and angular gyrus (Figure 5b). Area ‘9’ showed particularly robust co-activation with key hubs of the default network, such as anterior mPFC and posterior cingulate cortex (PCC), firmly placing this region in the default network despite its proximity to mid-DLPFC. In contrast, ‘IFG [L]’ showed a relatively distinct pattern, showing co-activation with portions of the fronto-parietal network– such as mid-DLPFC and pre-SMA. This pattern is consistent with the fact that left IFG’s contralateral homologue clustered with the fronto-parietal network and suggests this region may not be entirely dissociable from fronto-parietal clusters. Moreover, left IFG also showed stronger co-activation with posterior superior temporal sulcus– a key region implicated in semantic processing^15^ suggesting left IFG may also show a preference towards language topics.

**Figure 5.**
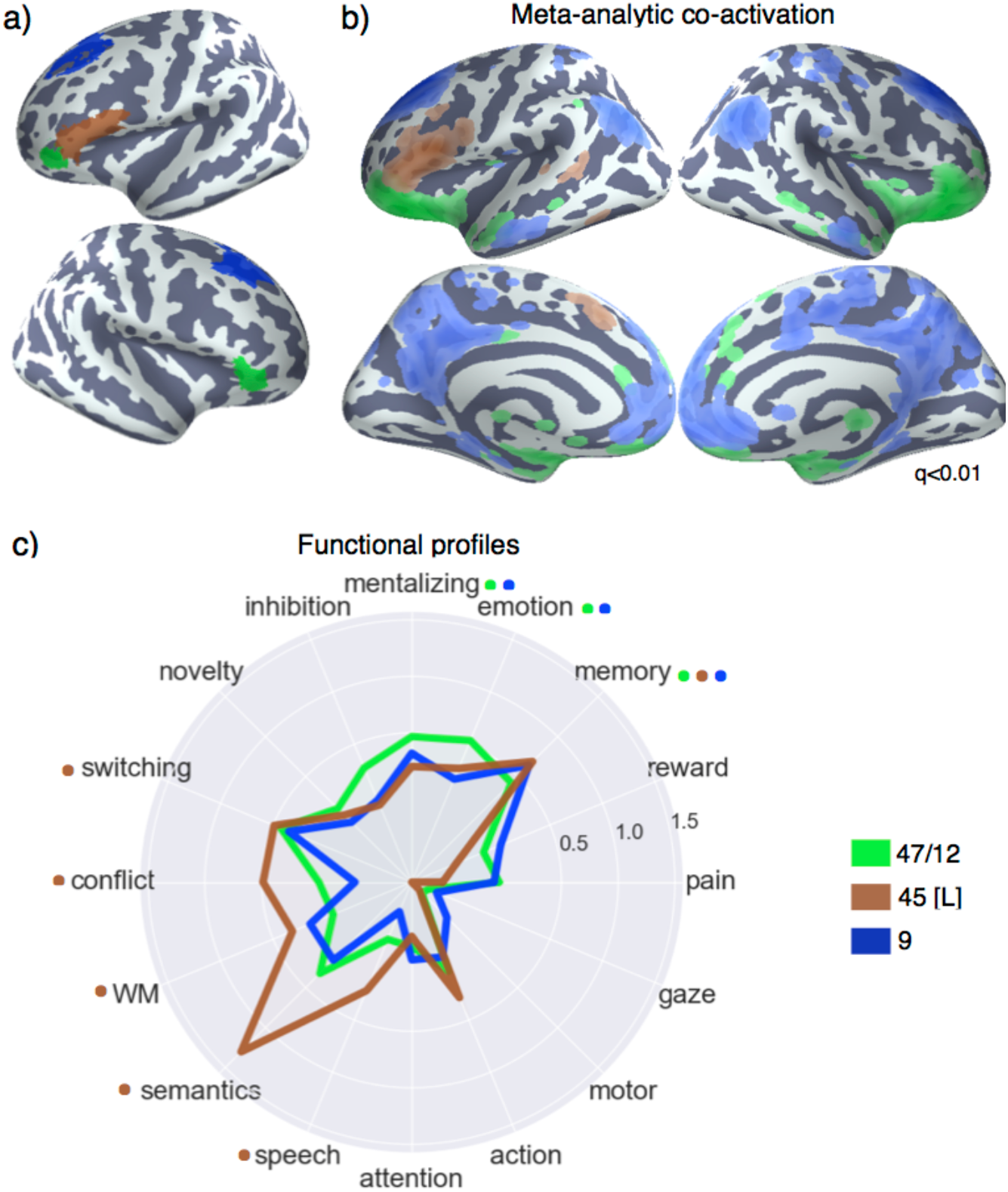
Lateral frontal regions of the default network. a) Individual clusters projected onto an inflated surface. b) Differences in coactivation between the three regions. Colored voxels activated more frequently in studies in the seed cluster of the same color was also active. c) Functional preference profiles for each cluster, revealing distinct psychological signatures for each subregion. Strength of association is measured in log oddsratio (LOR), and permutation-based significance is indicated next to each topic by color-coded dots corresponding to each region. Negative associations are indicated by the grey circle.

#### Meta-analytic functional preference

In contrast to clusters in the frontal-parietal network, clusters ‘47/12’ and ‘9’ showed no association with any executive processes– particularly notable for cluster ‘9’ due to its spatial proximity to mid fronto-parietal clusters (Figure 5c). Instead, clusters ‘47/12’ and ‘9’ were significantly associated with ‘mentalizing’, consistent with the hypothesis that these regions, as part of the dorsal medial subsystem of the default network, play an important role in conceptual processing and mentalizing^51^.

Distinct from other default network clusters, ‘45 [L]’ showed a significant association with various executive functions– further highlighting the distributed nature of executive processes across frontal regions. However, area ‘45 [L]’ was not associated with inhibition, suggesting inhibition is right-lateralized to some degree^52^. Furthermore, consistent with this region’s co-location with Broca’s area and co-activation with the superior temporal sulcus, area ‘45 [L]’ was significantly associated with ‘semantics’ and ‘speech’. Importantly, although it well-known Broca’s area is important for motor function in language, we did not find any association between area ‘45 [L]’ and motor topics; these results suggest Broca’s area is involved in the generation of speech motor plans, but not motor function more generally^53^. Moreover, ‘45 [L]’ was notable for its robust association with ‘semantic’ function– more so than any other region– consistent with the hypothesis that left IFG is a part of the brain’s ‘semantic’ system^15^.

Finally, consistent with the default network’s well-characterized involvement in memory^51^, all three LFC default clusters were robustly associated with ‘memory’ and ‘emotion’. This is consistent with a long line of evidence supporting the role of these regions in autobiographical, internally oriented cognition. Moreover, the left IFG is purported to play a key role in controlled memory retrieval^54,55^– a hypothesis supported by the joint association between executive processes and memory in this region. However, it is also notable that ‘memory’ was associated with many other clusters in the fronto-parietal network, suggesting memory processes are widely distributed across lateral frontal cortex.

### Somatosensory-motor network

We identified two LFC clusters in the somatosensory-motor network: dorsal and ventral lateral premotor cortex– ‘PMd’ and ‘PMv’, respectively (Figure 6a). Both clusters were located in dorsal BA 6^56^, although ‘PMd’ was slightly more anterior. As a result of its more posterior location, ‘PMv’ included several voxels in primary motor cortex, although the cluster was primarily located in pre-motor cortex.

**Figure 6.**
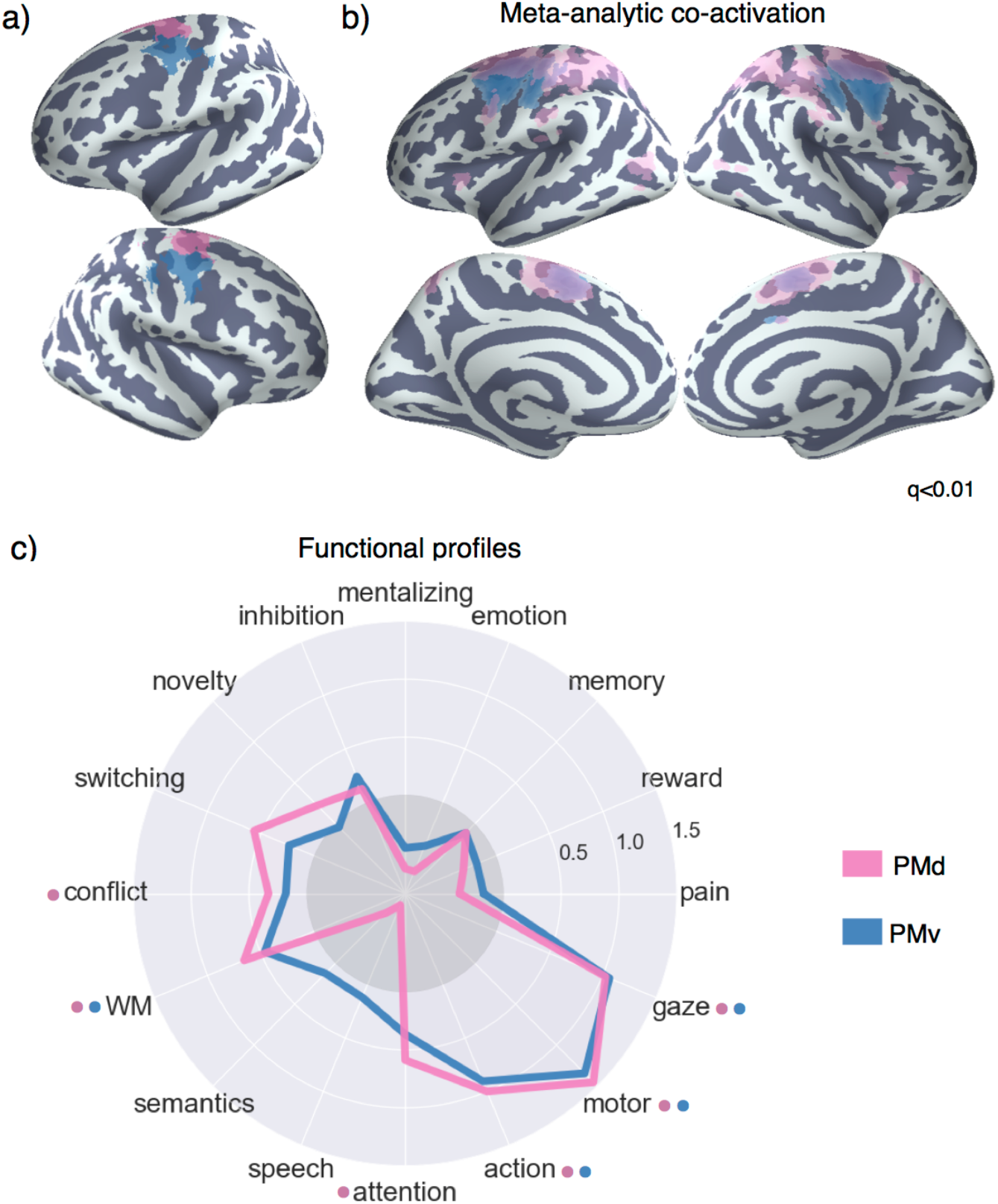
Meta-analysis of somatosensory clusters. a) Clusters projected onto an inflated surface b) Differences in co-activation between each cluster and the rest of LFC. Colored voxels activated more frequently in studies in which the seed cluster of the same color was also active. c) Functional preference profiles reveal distinct psychological signatures. Strength of association is measured in log odds-ratio (LOR), and permutation-based significance (q<0.05) is indicated next to each topic by color-coded dots corresponding to each cluster. Negative associations are indicated by the grey circle.

#### Meta-analytic co-activation

Both PMd and PMv showed greater co-activation with nearby voxels in the primary motor and somatosensory cortices, as well as SMA– regions important for the control of movement (Figure 6b). PMd, however, additionally showed greater co-activation with various regions implicated in executive function, such as lateral parietal cortex and the anterior insula– suggesting dorsal pre-motor cortex may engage a broader functional network in support of the cognitive control of motor actions.

#### Meta-analytic functional preference

The functional preference profiles of both premotor clusters suggest their primary functional role is in core aspects of motor function (Figure 6c). However, both of these the two clusters were also associated with higher-level motor planning (i.e. ‘action’) and working-memory, suggesting these regions are important for higher-level motoric control. Moreover, consistent with PMD’s stronger co-activation with regions previously associated with executive function, PMd was significantly associated with ‘conflict’ and ‘attention’ (although not significantly more so than PMv). Thus, although these two premotor clusters were most strongly associated with motor function, their function is not exclusively limited to low-level processes, and may require the recruitment of higher-level psychological processes for the execution of motor plans.

### Functional distance between clusters

**Figure 7.**
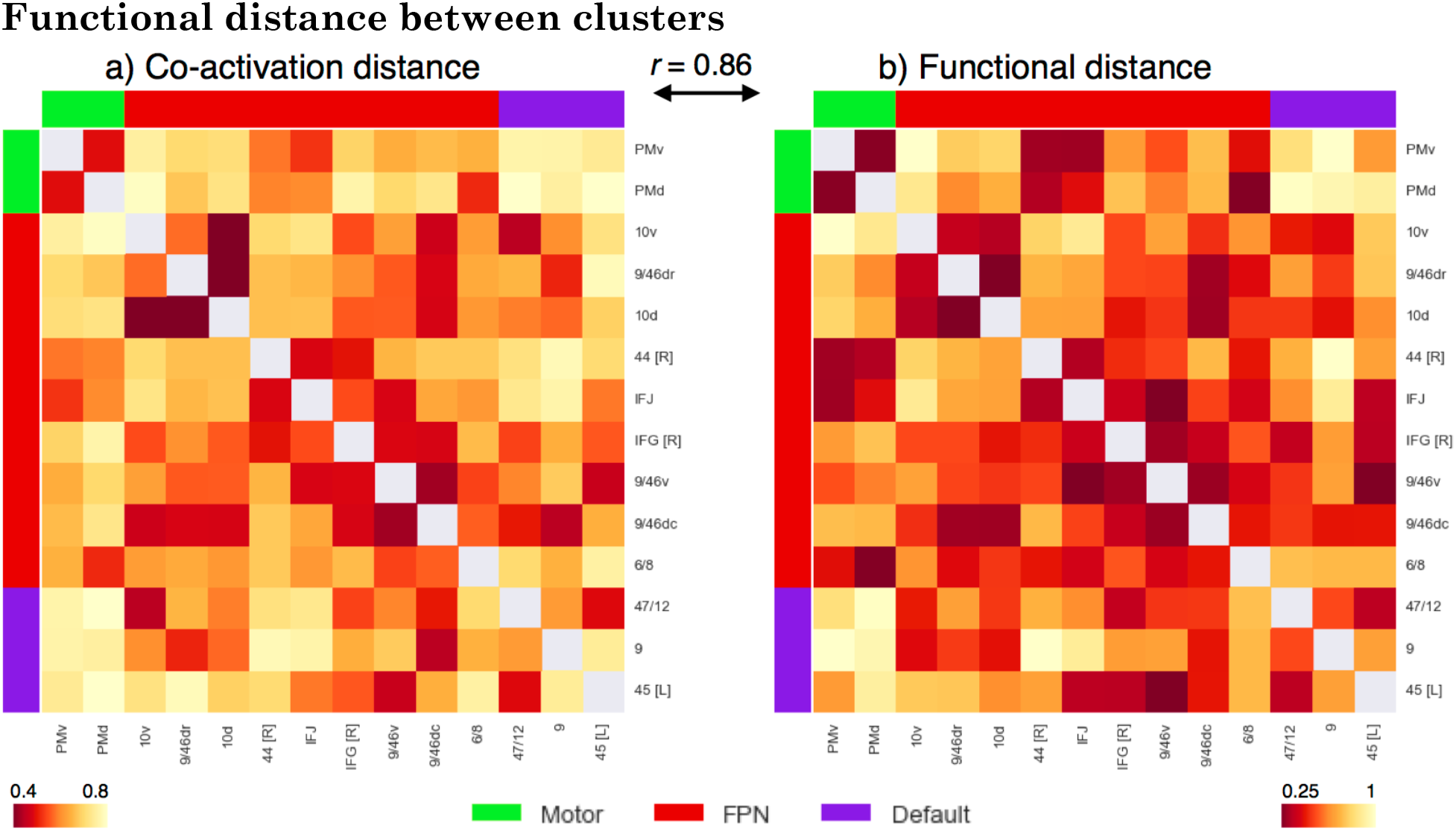
Co-activation and functional distance between LFC clusters. Pearson’s correlation distance between the 14 LFC clusters on the basis of meta-analytic (a) co-activation and (b) functional preference profiles. Although clusters within each network showed generally shorter distances to clusters in the same network than between networks, relatively high functional heterogeneity within each network was observed. The high similarities between these two distance matrices (r = 0.86, p < 0.001), suggests that the differences between regions observed in meta-analytic co-activation are generally accompanied by differences in functional preference profiles. Correlation distances range from 0 to 2, with 2 indicating perfect anti-correlation.

Finally, to examine the overall difference between regions, we computed the mean correlation distance between clusters on the basis meta-analytic co-activation (Figure 7a) and psychological preference profiles (Figure 7b). Supporting the network organization of these clusters, the distance between clusters in the same network was much shorter (co-activation: r=0.58, functional profiles, r=0.5) than the distance between clusters in different networks (co-activation: r=0.7, functional profiles, r=0.7) across both modalities. However, the distance between clusters in the same network was in certain cases relatively high. For example, clusters ‘45 [L]’ and ‘9’ in the default network (r = 0.77) and ‘44 [R]’ and ‘10v’ in the fronto-parietal network (r= 0.93) exhibited large functional distances, despite belonging to the same network. Thus, although large-scale networks likely represent a fundamental organizational structure in the brain– and distinct networks tend to support categorically different types psychological processes– our results suggest these networks are relatively heterogeneous. Finally, we also observed that the differences between regions based on meta-analytic co-activation were highly similar to those based on functional preference profiles (Pearson’s correlation r = 0.86), suggesting that clusters that show distinct meta-analytic co-activation generally exhibit distinct functional preference profiles.

## Discussion

In the present study, we applied data-driven methods to the largest meta-analytic database available to systematically map psychological states to discrete lateral frontal cortex anatomy. Importantly, we conducted our analyses broadly both with respect to anatomy– by focusing on the entirety of LFC– and function– by surveying a wide, representative range of psychological states– resulting in a relatively unbiased and comprehensive functional-anatomical mapping. Using co-activation-based hierarchical clustering, we identified 14 subregions in LFC organized into three whole-brain networks (fronto-parietal, default and sensorimotor). We then used multivariate classification to determine which psychological states best predicted activation in each region, resulting in dissociable psychological profiles for each subregion. In contrast with more modular models of LFC organization^57–59^, we observed a complex many-to-many mapping between individual regions and discrete psychological states, suggesting cognitive processes are supported in a distributed fashion by regions organized into whole-brain networks.

Consistent with the emerging view that the brain is composed of complex, distributed networks^18,32,33^, we found that individual regions within the same network exhibited relatively similar psychological profiles to each other. For example, all regions in the fronto-parietal network exhibited strong associations with executive functions, consistent with the hypothesis that the fronto-parietal network critical for flexible externally oriented behavior. In contrast, regions in different networks showed distinct psychological profiles from each other– despite occasionally high spatial proximity. For example, area ‘9’ of the default network, showed no significant association with any executive functions despite being positioned immediately dorsal to area ‘9/46’ of the fronto-parietal network. However, despite being relatively distant, areas ‘9’ and ‘47/12’ of the default network were both preferentially recruited by internally oriented processes such as ‘mentalizing’, ‘emotion’ and ‘memory’– a pattern consistent with a hypothesized role of the default network in self-generated conceptual processing^51^.

Although networks exhibited relatively robust dissociations, within each network we observed relatively low modularity, in contrast to more localizationist models. For example, sustained activity in DLPFC during working memory tasks has been hypothesized to reflect the active storage of working memory representations in domain-specific buffers^58^. However, we find that working memory recruits activity across a wide range of regions extending from posterior LFC to the lateral frontal pole. Moreover, many of these same regions that are preferentially recruited by working memory are similarly recruited by other executive functions, such as ‘conflict’ and ‘switching’, suggesting sustained activity in these regions supports domain-general processes required to flexibly guide behavior in support of the task goals^60^. These findings are consistent with a recent hypothesis that working memory is supported by the distributed reactivation of representations in parietal cortex, rather than isolated and modular maintenance in DLPFC^1,61^. In the same vein, updating task representations during task set switching has been hypothesized to preferentially recruit the inferior frontal junction^13,36^. However, we find that ‘switching’ preferentially recruits activity across a wide variety of LFC subregions as far rostral as the frontal pole, suggesting task-set switching is supported by distributed regions across the fronto-parietal network.

Importantly, although we observed relatively low functional specialization for individual regions across LFC, the present findings do not negate the idea that local neuron populations may be functionally specific. On the contrary, extensive neurophysiological data suggests association cortex contains overlapping neuron populations with distinct—and often highly specific—functional profiles^46–48^. Since subregions on the scale readily identified by fMRI likely exhibit aggregated activity across distinct neuron types, the relatively low modularity of these clusters should not be surprising. Rather, given that the distribution of distinct neuron populations likely varies across association cortex, one would expect individual regions to exhibit subtly varying associations to a wide range of psychological states.

Indeed, in the present study we observed substantial functional heterogeneity within each network and dissociable psychological profiles for regions within the same network. That is, although psychological states are not modularized into individual regions, the multivariate psychological profiles we generated for each region can be used to ascribe distinct roles for each region within the broader network. For instance, although all fronto-parietal regions were associated with various core executive functions, only IFJ showed additionally robust associations with high and low level motor function. Thus, it is plausible that IFJ may play an important role in biasing motoric representations in support of high-level goals represented in a distributed fashion throughout the network. In contrast, area 9/46v in mid-DLPFC was the region most strongly recruited by core executive processes, but showed no associations with ‘lower-level’ processes such as attention and motor function, suggesting this region may be more important for the biasing of abstract representations in more domain-specific regions of posterior cortex^62^.

Although the present results provide a comprehensive view into the functional organization of LFC, several challenges remain. More broadly, a difficult challenge in cognitive neuroscience is developing the appropriate psychological constructs that distinguish activity in related brain regions. Appropriately modeling the differences between nuanced psychological concepts is particularly difficult for large-scale meta-analyses, as there is no established ontology of psychological constructs, unlike in fields such as genetics^63^. In the present study, we used a data-driven set of topics derived from the abstracts of fMRI papers to represent major psychological phenomena. Although these topics are a major improvement on more simple term-based features, due to their data-driven nature they are likely to imperfectly capture psychological dimensions that are hypothesized to be important for differentiating regions. For example, in our set of 60 topics, only a single topic represented long-term memory function, and likely combined memory retrieval and autobiographical memory processes. Although the Neurosynth framework allows researchers to develop custom meta-analyses that can be used to test apriori predictions, the myriad of combinations in which studies can be combined is not conducive to determining the psychological dimensions that best differentiate brain activity.

The classification-based approach we employed is a step in the direction of quantifying the extent to which a given set of psychological features explains variability in brain activity. A promising future direction is to use classification based approaches and feature engineering to find the psychological dimensions that best differentiate patterns in activity between related regions, such as regions within a network. In combination with the adoption of standardized cognitive ontologies, such as the Cognitive Atlas^64^, such large-scale approaches should help the development of novel theories of functional brain organization. Moreover, given the limited quality of the summarized coordinate based data in Neurosynth^65^ the widespread sharing of richer statistical images in databases such as NeuroVault^66^ will greatly improve the fidelity of future meta-analyses.

In the present study, we used relatively unbiased data-driven methods to comprehensively map psychological states to individual regions in lateral frontal cortex. These regions were organized within large-scale whole-brain networks and shared functional properties with other regions in the same network. Moreover, we found that various specific psychological processes that have been previously hypothesized to map onto specific brain regions were widely distributed throughout lateral frontal cortex. Yet, we identified dissociable functional profiles for each subregion, suggesting that lateral frontal cortex supports a wide variety of psychological states through a mixture of network-level dynamics and moderate degree of functional specialization.

## Methods

### Dataset

We analyzed version 0.6 of the Neurosynth database^23^, a repository of 11,406 fMRI studies and over 410,000 activation peaks that span the full range of the published neuroimaging literature. Each observation contains the peak activations for all contrasts reported in a study’s table as well as the frequency of all of the words in the article abstract. A heuristic but relatively accurate approach is used to detect and convert reported coordinates to the standard MNI space. As such, all activations and subsequent analyses are in MNI152 coordinate space. The scikit-learn Python package^67^ was used for all machine learning analyses. Analyses were performed using the core Neurosynth python tools (https://github.com/neurosynth/neurosynth).

### Data and code availability

Code and data to replicate these analyses on any given brain region at any desired spatial granularity are available as a set of IPython Notebooks (https://github.com/adelavega/neurosynth-lfc).

### Lateral frontal cortex mask

To select clusters from whole-brain clustering solutions in lateral frontal cortex, we defined an LFC anatomical mask. Crucially, we only used this mask to select clusters that fell within this mask, and not to exclude individual voxels. First, we included voxels with a greater than 30% chance of falling in the frontal lobes according to the Montreal Neurological Institute structural probabilistic atlas and excluded medial voxels within 14mm of the midline. To focus on lateral frontal cortex, we excluded voxels that were exclusively located on the orbital surface– ensuring to include lateral orbitofrontal voxels– by removing voxels in the superior and medial orbital gyri according to the AAL atlas and voxels with a greater than 30% probability of falling in ‘Frontal Operculum Cortex’ in the Harvard-Oxford atlas. Finally, we also excluded far ventral voxels of OFC (Z < −14mm) that were not excluded using anatomical atlases.

### Co-activation clustering

Next, we clustered individual grey-matter cortical voxels across the whole brain based on their meta-analytic co-activation with the whole brain across studies in the database (Figure 1a). In order to avoid potentially biased or arbitrary cluster boundaries, we clustered the whole cortex and selected clusters for further analysis that fell within an anatomically defined LFC mask. Critically, we did not mask out voxels that were slightly outside of our mask– we either included or excluded entire clusters. This was particularly important for clusters near the edge of our LFC mask– as functional boundaries may not conform to anatomical boundaries– and at coarse clustering solutions– given the well-established finding that at least 4-5 whole-brain networks include voxels in lateral frontal cortex**32**. For whole-cortex clustering, we excluded voxels with less than 30% probability of falling in grey matter according to the Harvard-Oxford anatomical atlas and those with very low activation in the database (less than 100 studies per voxel). In general, Neurosynth’s activation mask (derived from the standard MNI152 template distributed with FSL) corresponded highly with probabilistic locations of cerebral cortex, with the exception of portions of dorsal precentral gyrus– which showed low activation although it was more than 50% likely to be in cerebral cortex.

We calculated the co-activation between each cortical voxel and every other voxel in the brain (including sub-cortex) by determining how correlated their activity was across studies. Activation in each voxel is represented as a binary vector of length 11,406 (the number of studies). A value of 1 indicated that the voxel fell within 10 mm of an activation focus reported in a particular study, and a value of 0 indicated that it did not. Because correlating the activation of every cortical voxel with every other voxel in the brain would result in a very large matrix (112,358 cortical voxels x 171,534 whole-brain voxels) that would be very computationally costly to cluster so as to identify distinct LFC regions. Hence, we reduced the dimensionality of the whole brain to 100 components using principal components analysis (PCA; the precise choice of number of components does not materially affect the reported results). Next, we computed the Pearson correlation distance between every voxel in the MFC mask with each whole-brain PCA component, resulting in a matrix that described the frequency with which each cortical voxel co-activated with the rest of the brain.

As an additional pre-processing step, we standardized each cortical voxel’s co-activation with other brain voxels to ensure clustering would be driven by relative differences in whole brain co-activation and not the overall activation rate of each voxel. That is, if two voxels co-activated with similar voxels across the brain, we should consider them to be relatively similar even if one of those voxels activates more frequently (and thus has slightly stronger correlations with all voxels). This adjustment was particularly important, as preliminary analyses indicated that regions with very high rates of activation (e.g. pre-SMA/mid-cingulate cortex) more readily clustered into multiple clusters with few voxels, reflecting base rates in activation, although differences in their functional associations were minimal. Indeed, preliminary analyses confirmed that standardizing the co-activation matrix alleviated this concern. At k = 70, the mean activation rate of each cluster showed no correlation with voxel size when Z-scoring was used (r=0.05), as compared to when the raw co-activation matrix was used (r = −0.65) at k = 70. Additionally, the range of cluster sizes was compressed, resulting in more evenly sized clusters. Cluster sizes ranged from 352 to 4546 voxels using the raw activation, compared to a range of 560 to 2862 voxels using standardized co-activation.

We applied hierarchical clustering with WarD’s linkage to the normalized co-activation matrix, resulting in a whole-brain linkage matrix. WarD’s clustering was selected as this algorithm is recommended as a good compromise between accuracy (e.g., fit to data) and reproducibility for clustering fMRI data^68^. However, this clustering algorithm is seldom used for whole-brain clustering because the computational time increases cubically [Θ(N^3^)] as a function of samples. We employed the fastcluster algorithm^69^—a package of libraries that enable efficient hierarchical clustering [Θ(N^2^)]—to achieve whole-brain clustering.

Since the optimality of a given clustering depends in large part on investigators’ goals, the preferred level of analysis, and the nature and dimensionality of the available data, identifying the ‘correct’ number of clusters is arguably an intractable problem^31^. However, in order to attempt to objectively guide the choice of choice of number of clusters to further analyze, we selected viable solutions using the silhouette score– a measure of within-cluster cohesion. Crucially, as we were specifically interested in the fit of the clustering to lateral frontal cortex, we only calculated the silhouette score with respect to voxels within our lateral frontal cortex mask. The silhouette coefficient was defined as (b – a) / max(a, b), where a is the mean intra-cluster distance and b is the distance between a sample and the nearest cluster of which the sample is not a part. Solutions that minimized the average distance between voxels within each cluster received a greater score. Once having chose two whole brain solutions, we extracted LFC clusters from with a substantial percentage of voxels within our apriori LFC mask. We varied the percentage of voxels within our LFC mask required to include a region across granularities with the objective maximizing coverage in LFC without including extraneous clusters with little presence in LFC. We arrived at 12% of voxels in a cluster within LFC at k=5 and 75% of voxels at k=70.

To understand the anatomical correspondence of the resulting clusters, we consulted a variety of anatomical and cytoarchitechtonic atlases. To locate each cluster anatomically, we used the probabilistic Harvard-Oxford atlas (H-O) that is packaged with FSL. We also visually compared the location of our clusters to the Petrides’ (2005) and Jülich micro-anatomical atlases included in FSL^56^. Regions were assigned names in accordance to Brodmann areas (BA) whenever clusters were sufficiently small to correspond to a single area (e.g. ‘area 9/46v’). Clusters were given functional names when they spanned multiple cytoarchitechtonic areas (e.g. IFJ) or multiple clusters spanned a single cytoarchitechtonic area (e.g. PMd & PMv). Note that although names were assigned to ease the discussion of these regions, we do not make strong claims of correspondence between functionally and anatomically defined regions, as we observed several discrepancies throughout LFC.

### Co-activation profiles

Next, we analyzed the differences in whole brain co-activation between the resulting clusters (Figure 1b) in order to understand the patterns of co-activation that differentiates these clusters. To highlight differences between clusters, we contrasted the co-activation of each cluster to the mean co-activation of the entire LFC. To do so, we performed a meta-analytic contrast between studies that activated a given cluster, and studies that activated a LFC mask composed of all clusters. The resulting images identify voxels with a greater probability of co-activating with the cluster of interest than with LFC on average. For example, voxels in blue in Figure 5b indicate voxels that are active more frequently in studies in which ‘area 9’ is active than in studies in which other LFC on average is active. We calculated p-values for each voxel using a two-way chi-square test between the two sets of studies and thresholded the co-activation images using the False Discovery Rate (q<0.01). The resulting images were binarized for display purposes and visualized using the pysurfer Python library.

### Topic modeling

Although term-based meta-analysis maps in Neurosynth closely resemble the results of manual meta-analyses of the same concepts, there is a high degree of redundancy between terms (e.g. ‘episodes’ and ‘episodic’), as well as potential ambiguity as to the meaning of an individual word out of context (e.g. ‘memory’ can indicate working memory or episodic memory). To remedy this problem, we employed a reduced semantic representation of the latent conceptual structure underlying the neuroimaging literature: a set of 60 topics derived using latent dirichlet allocation (LDA) topic-modeling30. This procedure was identical to that used in a previous study39, except for the use of a smaller number of topics and a much larger version of the Neurosynth database. The generative topic model derives 60 independent topics from the co-occurrence of all words in the abstracts of fMRI studies in the database. Each topic loads onto individual words to a varying extent, facilitating the interpretation of topics; for example, a working memory topic loads highest on the words “memory, WM, load”, while an episodic memory topic loads on “memory, retrieval, events”. Note that both topics highly load on the word “memory”, but the meaning of this word is disambiguated because it is contextualized by other words that strongly load onto that topic. Although the set of topics included 25 topics representing non-psychological phenomena– such as the nature of the subject population (e.g. gender, special populations) and methods (e.g., words such as “images”, “voxels”)—these topics were not explicitly excluded as they were rarely the strongest loading topics for any region. For all of our results, we focus on a set of 16 topics that strongly loaded onto lateral frontal cortex clusters (Table 1). These topics were obtained by determining the two strongest loading topics for each region.

**Table 1.** Topics most strongly associated with lateral frontal regions. Eight strongest loading words for each topic are listed, in descending order of association strength.

| Topic name | Top words |
| --- | --- |
| action | action actions motor goal mirror planning imitation execution |
| attention | attention attentional visual spatial search location orienting target |
| conflict | conflict interference incongruent stroop congruent selection competition color |
| emotion | emotional emotion regulation affective pictures emotions arousal affect |
| gaze | eye gaze eyes movements saccades target saccade visual |
| inhibition | inhibition inhibitory stop motor sustained nogo transient suppression |
| memory | memory retrieval encoding recognition episodic items recall words |
| mentalizing | social empathy moral person judgments mentalizing mental mind |
| motor | motor movement movements sensorimotor finger somatosensory sensory force |
| novelty | target targets novelty oddball distractor distractors deception mismatch |
| pain | pain stimulation somatosensory painful intensity sensory chronic noxious |
| reward | reward sleep anticipation monetary rewards motivation incentive loss |
| semantics | semantic words word lexical verbs abstract meaning verb |
| speech | speech auditory sounds sound perception voice acoustic listening |
| switching | switching rule executive switch rules flexibility shifting aggression |
| WM | memory working wm load verbal maintenance delay encoding |

### Meta-analytic functional preference profiles

We generated functional preference profiles by determining which psychological topics best predicted each cluster’s activity across fMRI studies (Figure 1c). First, we selected two sets of studies: studies that activated a given cluster– defined as activating at least 5% of voxels in the cluster– and studies that did not– defined as activating no voxels in the cluster. For each cluster, we trained a naive Bayes classifier to discriminate these two sets of studies based the loading of psychological topics onto individual studies. We chose naive Bayes because (i) we have previously had success applying this algorithm to Neurosynth data23; (ii) these algorithms perform well on many types of data, (iii) they require almost no tuning of parameters to achieve a high level of performance; and (iv) they produce highly interpretable solutions, in contrast to many other machine learning approaches (e.g., support vector machines or decision tree forests).

We trained models to predict whether or not fMRI studies activated each cluster, given the semantic content of the studies. In other words, if we know which psychological topics are mentioned in a study how well can we predict whether the study activates a specific region? We used 4-fold cross-validation for testing and calculated the mean score across all folds as the final measure of performance. We scored our models using the area under the curve of the receiver operating characteristic (AUC-ROC)– a summary metric of classification performance that takes into account both sensitivity and specificity. AUC-ROC was chosen because this measure is not detrimentally affected by unbalanced data^70^, which was important because each region varied in the ratio of studies that activated it to the studies that did not.

To generate functional preference profiles, we extracted from the naive Bayes models the log odds-ratio (LOR) of a topic being present in active studies versus inactive studies. The LOR was defined, for each region, as the log of the ratio between the probability of a given topic in active studies and the probability of the topic in inactive studies, for each region. LOR values above 0 indicate that a psychological topic is predictive of activation of a given region. To determine the statistical significance of these associations, we permuted the class labels and extracted the LOR for each topic 1000 times. This resulted in a null distribution of LOR for each topic and each cluster. Using this null distribution, we calculated p-values for each pairwise relationship between psychological concepts and regions, and reported associations significant after controlling for multiple comparisons using False Discovery Rate with q<0.01. Finally, to determine if certain topics showed greater preference for one cluster versus another, we conducted exploratory, post-hoc comparisons by determining if the 95% confidence intervals (CI) of the LOR of a specific topic for a one region overlapped with the 95% CI of the same topic in another region. We generated CIs using bootstrapping, sampling with replacement and recalculating log-odds ratios for each region 1000 times. A full reference figure of the loadings between topic and regions, including CIs, is available in Supplemental Figure 3. The ordering of the labels around the polar plot was determined using hierarchical clustering with average linkage, resulting in an order that concisely conveyed the functional differences between LFC’s subregions.

## Acknowledgments

R01MH096906 National Institutes of Health.

## Notes

Conflicts of Interest: The authors declare no competing financial interests.

## References

1. Miller, E. K. & Cohen, J. D. An integrative theory of prefrontal cortex function. Annu. Rev. Neurosci. 24, 167–202 (2001).

2. Miyake, A. et al. The unity and diversity of executive functions and their contributions to complex ‘Frontal Lobe’ tasks: a latent variable analysis. Cogn Psychol 41, 49–100 (2000).

3. Petrides, M. Lateral prefrontal cortex: architectonic and functional organization. Philosophical Transactions of the Royal Society B: Biological Sciences 360, 781–795 (2005).

4. Orr, J. M., Smolker, H. R. & Banich, M. T. Organization of the Human Frontal Pole Revealed by Large-Scale DTI-Based Connectivity: Implications for Control of Behavior. PLoS ONE 10, e0124797 (2015).

5. Neubert, F.-X., Mars, R. B., Sallet, J. & Rushworth, M. F. S. Connectivity reveals relationship of brain areas for reward-guided learning and decision making in human and monkey frontal cortex. Proceedings of the National Academy of Sciences 112, E2695–E2704 (2015).

6. Sallet, J. et al. The organization of dorsal frontal cortex in humans and macaques. J. Neurosci. 33, 12255–12274 (2013).

7. Kim, J.-H. et al. Defining functional SMA and pre-SMA subregions in human MFC using resting state fMRI: Functional connectivity-based parcellation method. NeuroImage 49, 2375–2386 (2010).

8. Goulas, A., Uylings, H. B. M. & Stiers, P. Unravelling the Intrinsic Functional Organization of the Human Lateral Frontal Cortex: A Parcellation Scheme Based on Resting State fMRI. Journal of Neuroscience 32, 10238–10252 (2012).

9. Eickhoff, S. B. et al. Assignment of functional activations to probabilistic cytoarchitectonic areas revisited. NeuroImage 36, 511–521 (2007).

10. Nee, D. E. et al. A Meta-analysis of Executive Components of Working Memory. Cereb. Cortex 23, 264–282 (2013).

11. Wager, T. D. & Smith, E. E. Neuroimaging studies of working memory: a meta-analysis. Cogn Affect Behav Neurosci 3, 255–274 (2003).

12. Nee, D. E., Wager, T. D. & Jonides, J. Interference resolution: insights from a meta-analysis of neuroimaging tasks. Cogn Affect Behav Neurosci 7, 1–17 (2007).

13. Derrfuss, J., Brass, M., Neumann, J. & Cramon, von, D. Y. Involvement of the inferior frontal junction in cognitive control: Meta-analyses of switching and Stroop studies. Hum. Brain Mapp. 25, 22–34 (2005).

14. Wager, T. D., Jonides, J. & Reading, S. Neuroimaging studies of shifting attention: a meta-analysis. NeuroImage 22, 1679–1693 (2004).

15. Binder, J. R., Desai, R. H., Graves, W. W. & Conant, L. L. Where Is the Semantic System? A Critical Review and Meta-Analysis of 120 Functional Neuroimaging Studies. Cereb. Cortex 19, 2767–2796 (2009).

16. Gilbert, S. J. et al. Functional specialization within rostral prefrontal cortex (area 10): a meta-analysis. Journal of Cognitive Neuroscience 18, 932–948 (2006).

17. Denny, B. T., Kober, H., Wager, T. D. & Ochsner, K. N. A meta-analysis of functional neuroimaging studies of self- and other judgments reveals a spatial gradient for mentalizing in medial prefrontal cortex. Journal of Cognitive Neuroscience 24, 1742–1752 (2012).

18. Petersen, S. E. & Sporns, O. Brain Networks and Cognitive Architectures. Neuron 88, 207–219 (2015).

19. Poldrack, R. A. Can cognitive processes be inferred from neuroimaging data? Trends in Cognitive Sciences 10, 59–63 (2006).

20. Dosenbach, N. U. F. et al. A core system for the implementation of task sets. Neuron 50, 799–812 (2006).

21. Duncan, J. The multiple-demand (MD) system of the primate brain: mental programs for intelligent behaviour. Trends in Cognitive Sciences 14, 172–179 (2010).

22. Nelson, S. M. et al. Role of the anterior insula in task-level control and focal attention. Brain Struct Funct 214, 669–680 (2010).

23. Yarkoni, T., Poldrack, R. A., Nichols, T. E., Van Essen, D. C. & Wager, T. D. Large-scale automated synthesis of human functional neuroimaging data. Nat. Methods 8, 665–670 (2011).

24. Toro, R., Fox, P. T. & Paus, T. Functional coactivation map of the human brain. Cereb. Cortex 18, 2553–2559 (2008).

25. Kober, H. & Wager, T. D. Meta?analysis of neuroimaging data. Wiley Interdisciplinary Reviews: Cognitive Science 1, 293–300 (2010).

26. De La Vega, A., Chang, L. J., Banich, M. T., Wager, T. D. & Yarkoni, T. Large-Scale Meta-Analysis of Human Medial Frontal Cortex Reveals Tripartite Functional Organization. J. Neurosci. 36, 6553–6562 (2016).

27. Pauli, W. M., O’Reilly, R. C., Yarkoni, T. & Wager, T. D. Regional specialization within the human striatum for diverse psychological functions. Proc. Natl. Acad. Sci. U.S.A. 113, 1907–1912 (2016).

28. Wager, T. D. et al. A Bayesian model of category-specific emotional brain responses. PLoS Comput Biol 11, e1004066 (2015).

29. Kober, H. et al. Functional grouping and cortical-subcortical interactions in emotion: a meta-analysis of neuroimaging studies. NeuroImage 42, 998–1031 (2008).

30. Blei, D. M., Ng, A. Y. & Jordan, M. I. Latent Dirichlet Allocation. Journal of Machine Learning Research 3, 993–1022 (2003).

31. Eickhoff, S. B., Thirion, B., Varoquaux, G. & Bzdok, D. Connectivity-based parcellation: Critique and implications. Hum. Brain Mapp. 36, 4771–4792 (2015).

32. Thomas Yeo, B. T. et al. The organization of the human cerebral cortex estimated by intrinsic functional connectivity. Journal of Neurophysiology 106, 1125–1165 (2011).

33. Power, J. D. et al. Functional network organization of the human brain. Neuron 72, 665–678 (2011).

34. Andrews-Hanna, J. R. The Brain's Default Network and Its Adaptive Role in Internal Mentation. The Neuroscientist 18, 251–270 (2012).

35. Paus, T. Location and function of the human frontal eye-field: a selective review. Neuropsychologia 34, 475–483 (1996).

36. Muhle-Karbe, P. S. et al. Co-Activation-Based Parcellation of the Lateral Prefrontal Cortex Delineates the Inferior Frontal Junction Area. Cereb. Cortex 26, 2225–2241 (2016).

37. Bludau, S. et al. Cytoarchitecture, probability maps and functions of the human frontal pole. NeuroImage 93 Pt 2, 260–275 (2014).

38. Chang, L. J., Yarkoni, T., Khaw, M. W. & Sanfey, A. G. Decoding the Role of the Insula in Human Cognition: Functional Parcellation and Large-Scale Reverse Inference. Cereb. Cortex 23, 739–749 (2013).

39. Poldrack, R. A. et al. Discovering Relations Between Mind, Brain, and Mental Disorders Using Topic Mapping. PLoS Comput Biol 8, e1002707–14 (2012).

40. Nee, D. E. & Brown, J. W. Rostral–caudal gradients of abstraction revealed by multi-variate pattern analysis of working memory. NeuroImage 63, 1285– 1294 (2012).

41. Mackey, W. E., Devinsky, O., Doyle, W. K., Meager, M. R. & Curtis, C. E. Human Dorsolateral Prefrontal Cortex Is Not Necessary for Spatial Working Memory. Journal of Neuroscience 36, 2847–2856 (2016).

42. De Baene, W., Albers, A. M. & Brass, M. The what and how components of cognitive control. NeuroImage 63, 203–211 (2012).

43. Wager, T. D., Davidson, M. L., Hughes, B. L., Lindquist, M. A. & Ochsner, K. N. Prefrontal-subcortical pathways mediating successful emotion regulation. Neuron 59, 1037–1050 (2008).

44. Woo, C.-W. et al. Separate neural representations for physical pain and social rejection. Nature Communications 5, 5380 (2014).

45. Chatham, C. H. et al. Cognitive control reflects context monitoring, not motoric stopping, in response inhibition. PLoS ONE 7, e31546 (2012).

46. Kvitsiani, D. et al. Distinct behavioural and network correlates of two interneuron types in prefrontal cortex. Nature 498, 363–366 (2013).

47. Tye, K. M. & Deisseroth, K. Optogenetic investigation of neural circuits underlying brain disease in animal models. Nat Rev Neurosci 13, 251–266 (2012).

48. Xiu, J. et al. Visualizing an emotional valence map in the limbic forebrain by TAI-FISH. Nat Neurosci 17, 1552–1559 (2014).

49. Petrides, M. & Pandya, D. N. Projections to the frontal cortex from the posterior parietal region in the rhesus monkey. J. Comp. Neurol. 228, 105– 116 (1984).

50. Cavada, C. & Goldman-Rakic, P. S. Posterior parietal cortex in rhesus monkey: I. Parcellation of areas based on distinctive limbic and sensory corticocortical connections. J. Comp. Neurol. 287, 393–421 (1989).

51. Andrews-Hanna, J. R., Smallwood, J. & Spreng, R. N. The default network and self-generated thought: component processes, dynamic control, and clinical relevance. Ann. N.Y. Acad. Sci. 1316, 29–52 (2014).

52. Aron, A. R., Robbins, T. W. & Poldrack, R. A. Inhibition and the right inferior frontal cortex: one decade on. Trends in Cognitive Sciences 18, 177–185 (2014).

53. Flinker, A. et al. Redefining the role of Broca's area in speech. Proc. Natl. Acad. Sci. U.S.A. 112, 2871–2875 (2015).

54. Badre, D. & Wagner, A. D. Left ventrolateral prefrontal cortex and the cognitive control of memory. Neuropsychologia 45, 2883–2901 (2007).

55. Snyder, H. R., Banich, M. T. & Munakata, Y. Choosing our words: retrieval and selection processes recruit shared neural substrates in left ventrolateral prefrontal cortex. Journal of Cognitive Neuroscience 23, 3470–3482 (2011).

56. Eickhoff, S. B. et al. Assignment of functional activations to probabilistic cytoarchitectonic areas revisited. NeuroImage 36, 511–521 (2007).

57. Bertolero, M. A., Yeo, B. T. T. & D’Esposito, M. The modular and integrative functional architecture of the human brain. Proc. Natl. Acad. Sci. U.S.A. 112, E6798–807 (2015).

58. Baddeley, A. Working memory: looking back and looking forward. Nat Rev Neurosci 4, 829–839 (2003).

59. Fodor, J. A. The modularity of mind: An essay on faculty psycholgy. (Bradford Bks, 1983).

60. Curtis, C. E. & Lee, D. Beyond working memory: the role of persistent activity in decision making. Trends in Cognitive Sciences 14, 216–222 (2010).

61. Postle, B. R. in 1–14 (2016).

62. Badre, D. Cognitive control, hierarchy, and the rostro–caudal organization of the frontal lobes. Trends in Cognitive Sciences 12, 193–200 (2008).

63. Ashburner, M. et al. Gene ontology: tool for the unification of biology. The Gene Ontology Consortium. Nat. Genet. 25, 25–29 (2000).

64. Poldrack, R. A. et al. The cognitive atlas: toward a knowledge foundation for cognitive neuroscience. Front Neuroinform 5, 17 (2011).

65. Salimi-Khorshidi, G., Smith, S. M., Keltner, J. R., Wager, T. D. & Nichols, T. E. Meta-analysis of neuroimaging data: a comparison of image-based and coordinate-based pooling of studies. NeuroImage 45, 810–823 (2009).

66. Gorgolewski, K. J. et al. NeuroVault.org: a web-based repository for collecting and sharing unthresholded statistical maps of the human brain. Front Neuroinform 9, 8 (2015).

67. Pedregosa, F. et al. Scikit-learn: Machine Learning in Python. Journal of Machine Learning Research 12, 2825–2830 (2011).

68. Thirion, B., Varoquaux, G., Dohmatob, E. & Poline, J.-B. Which fMRI clustering gives good brain parcellations? Front. Neurosci. 8, 169 (2014).

69. Müllner, D. fastcluster: Fast hierarchical, agglomerative clustering routines for R and Python. Journal of Statistical Software (2013).

70. Jeni, L. A., Cohn, J. F. & La Torre, De, F. Facing Imbalanced Data– Recommendations for the Use of Performance Metrics. in 2013, 245–251 (IEEE, 2013).

